# Two phosphoglucomutase paralogs regulate triggered secretion of the *Toxoplasma* micronemes

**DOI:** 10.1101/157743

**Authors:** Sudeshna Saha, Bradley I. Coleman, Tiffany Sansom, Rashmi Dubey, Ira J. Blader, Marc-Jan Gubbels

**Affiliations:** Department of Biology, Boston College, Chestnut Hill, MA 02467, USA; Department of Microbiology and Immunology, University at Buffalo School of Medicine, Buffalo, NY 14214, USA

**Keywords:** *Toxoplasma*, micronemes, calcium, parafusin, phosphoglucomutase

## Abstract

Parafusin is a phosphoglucomutase (PGM) paralog that acts as a signaling scaffold protein in calcium mediated exocytosis across many eukaryotes. In *Toxoplasma gondii* the parafusin related protein 1 (PRP1) has been associated in indirect and heterologous studies with the regulated exocytosis of the micronemes, which are required for successful host cell invasion and egress. Here we directly assessed the role of PRP1 by deleting the gene from the parasite. We observed a specific defect in microneme secretion in response to high Ca^2+^ fluxes, but not to phosphatidic acid fluxes controlling microneme release. We observed no defect in constitutive microneme secretion which was sufficient to support completion of the lytic cycle. Furthermore, deletion of the other PGM in *Toxoplasma*, PGM2, as well as the double PRP1/PGM2 deletion resulted in a similar phenotype. This suggests a functional interaction between these two genes. Strikingly, tachyzoites without both paralogs are completely viable *in vitro* and during acute mice infections. This indicates that PGM activity is neither required for glycolysis. In conclusion, the PRP1-PGM2 pair is required for a burst in microneme secretion upon high Ca^2+^ fluxes, but this burst is not essential to complete the lytic cycle of the parasite.

**Plain Language Summary:** Calcium mediated control of microneme secretion is essential for host cell invasion and egress of *Toxoplasma gondii*. Here it is shown that the two phosphoglucomutases in *Toxoplasma* both function in the translation of a spike in calcium into a burst in microneme secretion.

## Introduction

The apicomplexan parasite *Toxoplasma gondii* has infected 1 in every 3 humans globally but most acute infections pass with mild or no symptoms (Montoya & Liesenfeld, 2004). However, severe disease occurs congenitally in pregnant women (Remington *et al.*, 1995) and in immunocompromised patients (i.e. AIDS patients) where reactivation of a chronic infection can result in life-threatening encephalitis or myocarditis (Weiss & Dubey, 2009). The clinical manifestation results from the active invasion, intracellular replication and egress required by the lytic cycle. As such host cell invasion is a critical event in pathogenesis (Blader *et al.*, 2015). Unlike some other intracellular pathogens, host cell invasion by *Toxoplasma* is a parasite-directed event requiring parasite motility and sequential secretion of three different secretory organelles (Baum *et al.*, 2008, Meissner *et al.*, 2013). The secretion of the first organelle, the micronemes, is regulated by intracellular calcium (Ca^2+^) fluxes (Lourido & Moreno, 2015, Wetzel *et al.*, 2004, Sidik *et al.*, 2016a), however the molecular mechanism underlying this control, especially the late events facilitating exocytosis, is still incompletely understood.

The phosphoglucomutase (PGM) family comprises glycolytic enzymes that interconvert glucose-1-phosphate and glucose-6-phosphate. However, a PGM ortholog also known as parafusin or PFUS is present in many different organisms including ciliates, yeast and humans that functions in Ca^2+^ signaling (Levin *et al.*, 1999, Andersen *et al.*, 1994, Subramanian *et al.*, 1994, Satir *et al.*, 1989, Satir *et al.*, 2015). The *Toxoplasma* genome encodes two PGM paralogs: PGM1 (TGME49_285980), also referred to as parafusin-related protein 1 (PRP1), and PGM2 (TGME49_318580) (Imada *et al.*, 2010). In the ciliate *Paramecium* PFUS has been associated with Ca^2+^-mediated exocytosis of its dense core secretory vesicles (DCSVs) (Gilligan & Satir, 1982, Plattner & Kissmehl, 2005). In *Paramecium* PFUS associates with the DCSVs acting as scaffold with a direct role in membrane fusion (Zhao & Satir, 1998). Furthermore, PFUS also has a role in DCSV assembly in both *Paramecium* (Liu *et al.*, 2011) and *Tetrahymena* (Chilcoat & Turkewitz, 1997) and is hypothesized to impact the local rather than systemic release of the matured DCSVs (Plattner & Kissmehl, 2005). In ciliates PFUS is dynamically phosphorylated in a Ca^2+^-dependent fashion related to Ca^2+^-dependent DCSV exocytosis (Subramanian & Satir, 1992, Wyroba *et al.*, 1995, Satir *et al.*, 1990, Matthiesen *et al.*, 2003): in its phosphorylated state PFUS associates with the vesicles closest to the plasma membrane, priming them for secretion; however for actual membrane fusion to take place PFUS has to be dephosphorylated through calcineurin (Plattner & Kissmehl, 2005). More recently,additional roles for PFUS have been uncovered as it was found to be associated with the base of cilia and nucleus in ciliates as well as mammalian cells (Satir et al., 2015) whereas in yeast PFUS has been associated with regulating Ca^2+^ homeostasis (Fu *et al.*, 2000). Germane to *Toxoplasma*, the ortholog PRP1 has been shown to localize to the most apical micronemes, its phosphorylation status is Ca^2+^-dependent, and through heterologous evaluation of PRP1 in *Paramecium*, an orthologous function in Ca^2+^-mediated microneme secretion has been proposed (Matthiesen *et al.*, 2001, Matthiesen et al., 2003).

Here we directly evaluated the role of PRP1 and PGM2 through gene deletions in *Toxoplasma*. We show that both paralogs are required for microneme secretion in response to high Ca^2+^ spikes, which occur during host cell invasion and egress. However, both paralogs are expendable for constitutive microneme secretion under conditions mimicking gliding motility between host cells. Strikingly, tachyzoites without both paralogs are completely viable *in vitro* as well as during acute mice infections. These data suggest that the *Toxoplasma* PGMs are dispensable for Ca^2+^-dependent exocytosis and glycolysis during the lytic cycle.

## Results

### PRP1 is conserved across the coccidia

To establish whether PRP1 has a universal role in exocytosis across the Apicomplexa we first explored the conservation and phylogeny of PGMs in the Apicomplexan and their closely related free-living relatives, the Chromerids. Both TgPGM1 and TgPGM2 were compared to the validated PGM1/parafusins in ciliates and the crown eukaryotes (Fig 1). Consistent with the observation that PGM gene duplication occurred before the evolution of the eukaryotic cell (Satir et al., 2015), these results show that all PGM1 sequences cluster together and are uniquely distinct from the PGM2 sequences. The conservation of both PGM1 and PGM2 is very different for different apicomplexan sub-groups. Although *Toxoplasma* and the other cyst-forming coccidia (*Hammondia hammondi, Neospora caninum*, and *Sarcocystis neurona*) have both PGM1 and PGM2, the non-cyst forming coccidia, i.e. *Eimeria* spp., only encode one PGM1 ortholog. In the genus *Plasmodium*, *P. falciparum* only encodes PGM2 but not *P. chabaudi* and *P. berghei* contain both whereas *Cryptosporidium* spp and the related *Gregarina niphandrodes* only encode PGM1; in fact, *Cr. parvum* and *Cr. muris* encode two slightly different PGM1 isotypes encoded in tandem in the genome. We did not find any annotated PGM in the genome of *Theileria annulata.* On the other hand, both Chromerids (*Chromera Velia* and *Vitrella brassicaformis*) contain both PGM1 and PGM2. The ciliates *Paramecium tetraurelia* and *Tetrahymena thermophile* only contain PGM1. Overall, we interpret the data to mean that the common ancestor of the Alveolata contained both PGM1 and PGM2. This is still seen in some extant lineages like *Toxoplasma*, but in other lineages either one has been lost. Hence, since *Toxoplasma* contains both isoforms, it is an ideal organism to study the contribution of these unique glycolytic enzymes either to metabolism and/or Ca^2+^-dependent exocytosis.

**Figure 1:**
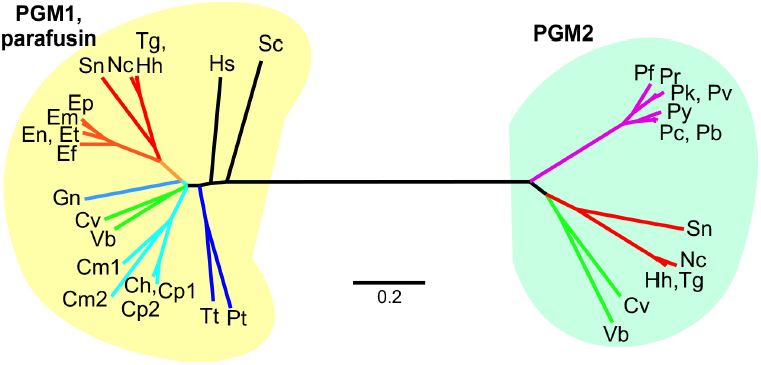
Phylogeny of PGMs in the Apicomplexa and their relatives. Unrooted tree based on alignment of the following sequences: *Toxoplasma gondii*: TGME49_285980 (TgPGM1), TGME49_318580 (TgPGM2); *Neospora canimun*:A0A0F7UAD1 (PGM1: NCLIV_014450 has wrong first exon), NCLIV_010960 (NcPGM2),*Hammondia hammondi*: HHA_285980 (HhGPM1), HHA_318580 (HhGPM2); *Sarcocystis neurona*: SN3_00900660 (SnPGM1), SN3_01700255 (SnPGM2); *Eimeria maxima*: EMWEY_00024400 (EmPGM1); *Eimeria praecox*: EPH_0036200 (EpPGM1); *Eimeria falciformis*: EfaB_MINUS_40637.g2669_1 (EfPGM1); *Eimeria tenella*: ETH_00002785 (EtPGM1); *Eimeria necatrix*: ENH_00082780 (EnPGM1); *Eimeria brunetti*: EBH_0076610 (EbPGM1); *Plasmodium falciparum*: PF3D7_1012500 (PfGGM2); *Plasmodium reichenowi*: PRCDC_1011900 (PrGGM2); *Plasmodium chabaudi*: PCHAS_1211600 (PcPGM2); *Plasmodium berghei*: PBANKA_1210900 (PbPGM2); *Plasmodium yoelii*: PY17X_1214100 (PyPGM2); *Plasmodium knowlesi*: PKNH_0812300 (PkPGM2); *Plasmodium vivax*: PVX_094845 (PvPGM2); *Gregarina niphandrodes*: GNI_111250(GnPGM1); *Cryptosporidium parvum*: cgd2_3270 (CpPGM1.1), cgd2_3260 (CpPGM1.2); *Cryptosporidium muris*: CMU_003300 (CmPGM1.1), CMU_003310 (CmPGM1.2); *Cryptosporidium hominis*: Chro.20343 (ChPGM1); *Chromera velia*: Cvel_11076 (CvPGM1), Cvel_22350 (CvPGM2); *Vitrella brassicaformis*: Vbra_3443 (VbPGM1), Vbra_19870 (VbPGM2). *Paramecium tetraurelia*: 1KFI_A (PtGPGM1); *Tetrahymena thermophile*: AAB97159.1 (TtPGM1); *Saccharomyces cerevisiae*: CAA89741.1 (ScPGM1); *Homo sapiens*: AAA60080.1 (HsPGM1). Branch colors reflect species relation with the cyst forming coccidia in red, non-cyst forming coccidia in orange, *Plasmodium* spp. in purple, *Cryptosporidium* spp. in light blue, gregarines in medium blue, ciliates in dark blue and the chromerids in green.

### PRP1 is not required for completion of the lytic cycle

The first question we addressed is whether PRP1 is essential. We generated a direct PRP1-KO parasite line by replacing the PRP1 gene with an HXGPRT selectable marker (Fig S1). To validate the loss of protein expression we generated a specific polyclonal antiserum against amino acids 446 to 637 in the C-terminal protein region of PRP1, which is not shared with PGM2 (Fig 2A). PRP1 is undetectable in the direct KO line (ΔPRP1). This result immediately indicated that PRP1 is not required to complete the lytic cycle. However, it is possible that the ΔPRP1 line has a growth disadvantage. We therefore performed plaque assays and compared plaque number and size to the parent line (Fig 2B). We observed similar plaque number for the parent and ΔPRP1 lines but a minor yet significant fitness advantage for the parasites lacking PRP1, as determined by total area of the plaques. Since PRP1 was first described for its involvement in microneme secretion we tested two essential processes relying on microneme secretion: host cell egress and invasion. We tested both the Ca^2+^-dependent and phosphatidic acid (PA)-dependent legs underlying microneme secretion (Bullen *et al.*, 2016) by titrating in A23187 Ca2+-ionophore and propranolol to activate diacylglycerol kinase 1 (DGK1) to raise the PA concentration, respectively. With neither treatment did we observe a difference in egress efficiency compared to the RHΔku80 parent line, suggesting that micronemes are secreted sufficiently to support egress (Fig 2C). We neither observed a difference in invasion efficiencies, indicating that secretion of the rhoptries is also not affected by the absence of PRP1 (Fig 2D). We more directly monitored rhoptry secretion by detecting phosphorylated STAT3 (P-STAT3) accumulating in in the nucleus of the infected host cell as STAT3 is phosphorylated by the rhoptry protein ROP16 (Saeij *et al.*, 2007). We did not observe a difference in accumulation of P-STAT3 between parent and ΔPRP1 lines supporting that rhoptries are secreted normally (Fig S2). Based on these results we conclude that PRP1 is not essential for the secretion of micronemes and rhoptries and is not required for completing the *Toxoplasma* lytic cycle *in vitro*.

**Figure 2.**
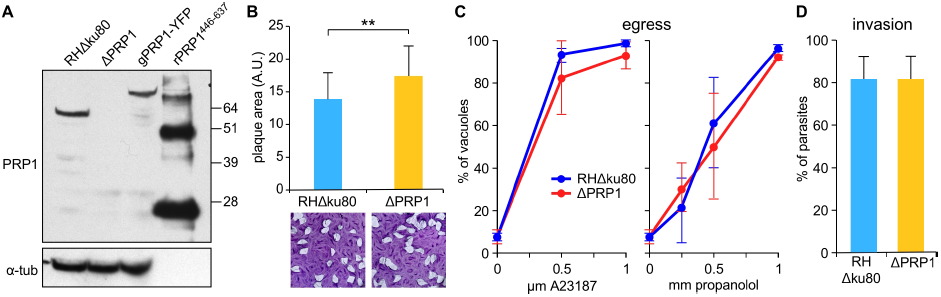
PRP1 antiserum validation and global phenotype analysis of ΔPRP1 parasites. **A.**Parental (RHΔku80), ΔPRP1 and endogenously YFP-tagged PRP1 (gPRP1-YFP) parasites were lysed and analyzed in a western blot using α-PRP1 antibody. The 27 kDa shift in the molecular weight resulting from the YFP tag can be observed in gPRP1-YFP lysate. rPRP1^446^-^637^ represents the His6-tagged recombinant protein used to generate the αPRP1 antiserum. α-tubulin antibody was used as loading control of the parasite lysate. **B.**Bottom: Representative image of plaque assay performed with ΔPRP1 and parental RHΔ*ku80* parasites grown undisturbed on a host cell monolayer for seven days.Top: Quantification of the plaque size following digital scanning of the plaque assays. A.U. arbitrary units. Counted 30 plaques per sample, n=4. Mean ± s.e.m.; two-tailed *t*-test: [**], *P* <0.001. **C.**ΔPRP1 and parent parasites were triggered to egress by the Ca^2+^ ionophore A23187 or propranolol to activate DGK1 and raise the PA concentration for five minutes at 37°C. For all samples, egress following stimulation is expressed as the percent egressed vacuoles of the total vacuoles observed. n=3; mean ±s.d. **D.** The invasion efficiency was assessed using the red-green assay with freshly lysed ΔPRP1 and parental RHΔku80 parasites. Total numbers of intracellular and extracellular parasites per microscopy field were counted and the intracellular parasites expressed relative to the total parasite. At least 150 parasites from three random fields per sample were scored. n=3; mean ±s.d.

### Loss of PRP1 reduces Ca^2+^-induced microneme secretion

To assess whether the absence of PRP1 more subtly affected signaling events leading to microneme secretion we used pharmacological triggers (secretagogues) acting on distinct steps in the signal transduction pathway. The current understanding of the signaling pathway is that protein kinase G (PKG) acts upstream of phosphoinositide phospholipase C (PI-PLC), which subsequently results in the formation of inositol triphosphate (IP_3_) and diacylglycerol (DAG) (Lourido & Moreno, 2015). Here the pathway bifurcates as IP_3_ leads to the release of Ca^2+^ from the endoplasmic reticulum, which is relayed by calcium-dependent protein kinases (CDPKs), whereas DAG is converted to PA, which is directly sensed by the micronemes to promote their secretion (Bullen *et al.*, 2016). We used zaprinast to activate PKG (Donald & Liberator, 2002, Lourido *et al.*, 2012), ethanol to trigger PI-PLC (Carruthers *et al.*, 1999), A23187 to mimic the high Ca^2+^ trigger (Carruthers & Sibley, 1999, Lourido *et al.*, 2010), and propranolol to activate DGK1 which raises the PA concentration and engages the sensor APH (acylated pleckstrin-homology domain-containing protein) on the micronemes (Jacot *et al.*, 2016, Bullen et al., 2016). High spikes in cytoplasmic Ca^2+^ concentration have been reported to coincide with egress and invasion whereas an intermediate elevated cytoplasmic Ca^2+^ concentration accompanies gliding motility between host cells (Sidik et al., 2016a). Hence, we determined the level of so-called ‘constitutive’ microneme secretion in absence of pharmacological secretagogues measured over one hour, reflecting the basal level of microneme secretion during extracellular gliding (Wetzel et al., 2004). Secretion was assessed through the release of proteolytically cleaved microneme protein, MIC2, in the supernatant of extracellular parasites. We observed robust constitutive secretion regardless of the presence or absence of PRP1 (Fig 3A). Even the activation of signaling events in the DAG leg of the pathway and further upstream (resulting from propranolol, ethanol, or zaprinast inductions) did not result in any dramatic change of MIC2 release, except for a minor but notable reduction of ethanol-triggered secretion in absence of PRP1. However, the ΔPRP1 parasites were less responsive to the Ca^2+^ branch of the signaling pathway triggered by Ca^2+^ ionophore A23187, which sharply reduced the amount of released MIC2. Thus, the role of PRP1 in microneme secretion is exclusive to the Ca^2+^-dependent signal transduction pathway and does not act on the PA-mediated branch contributing to microneme secretion.

**Figure 3.**
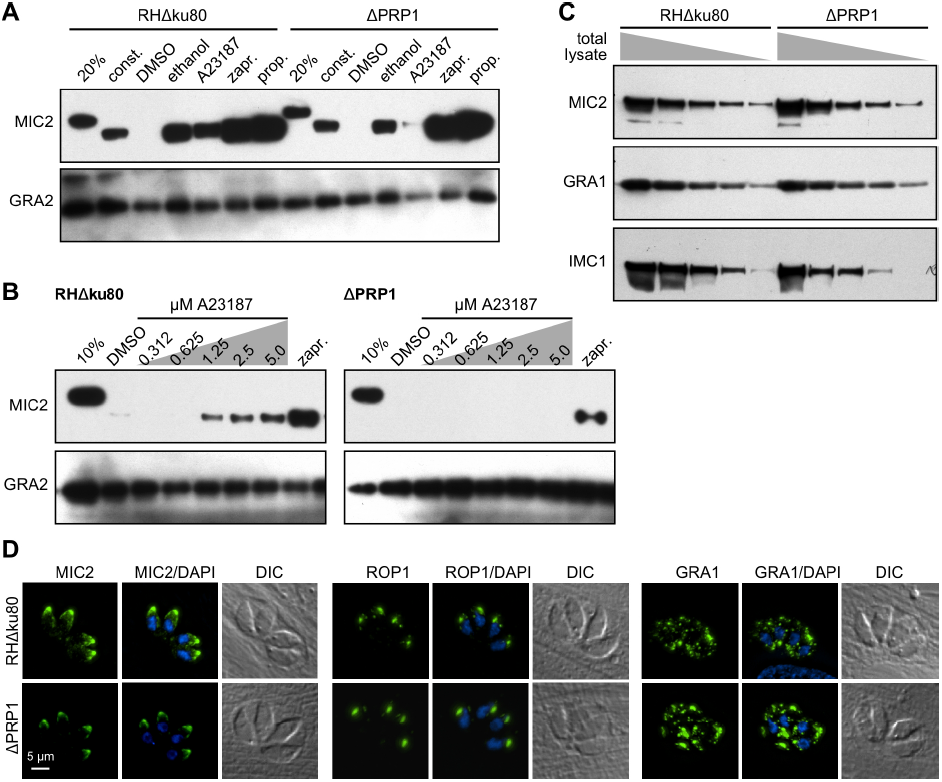
Absence of PRP1 disrupts high Ca^2+^-trigger induced microneme secretion in tachyzoites. **A.** Representative western blot image of the microneme secretion assay performed with ΔPRP1 and parent parasites. The secretion of the microneme protein MIC2 was used as the marker. 20% represents the non-secreted total protein lysate from 20% of the total parasites used in the secretion assay. Extracellular tachyzoites were treated with 0.25%(v/v) ethanol, 1.25 μ M A23187, 500 μ M zaprinast (zapr), 500 μ M propranolol (prop) or DMSO as control for five minutes at 37°C. For constitutive (const.) secretion, extracellular tachyzoites were allowed to release the protein for one hour in absence of pharmacological trigger. Proteolytic processing of the secreted microneme protein can be seen as shift in the MIC2 band. Dense granule protein GRA2 was used as a control for microneme- and Ca^2+^-independent secretion. **B.**Titration of the Ca^2+^ ionophore A23187 used for triggering microneme secretion in ΔPRP1 and RHΔku80 parent parasites. Extracellular tachyzoites were treated with the indicated concentrations of A23187 and zaprinast as control for five minutes at 37°C. 10% represents the non-secreted total protein lysate from 10% of the total parasites used in the secretion assay. **C.**Representative blot to assess the relative abundance of MIC2 protein in ΔPRP1 and RHΔku80 parent parasites lysates. Total parasite lysates were serially diluted and blots probed with the antisera as indicated. GRA1 (dense granules) and IMC1 (cytoskeleton) were used as loading control markers. **D.**Immunofluorescence displaying morphology of the secretory organelles in ΔPRP1 and RHΔku80 parent parasites. MIC2: micronemes; ROP1: rhoptries; GRA1 dense granules (all in green). DAPI marks DNA.

To differentiate whether the loss of the high Ca^2+^ response is the result of an overall deficiency or of a reduced sensitivity we titered the amount of A23187 in the microneme secretion assay (Fig 3B). Compared to wild type parasites that secrete MIC2 at A23187 concentrations as low as 1.25 μM and reached saturation at 2.5 μ M, ΔPRP1 mutants never display any secretion even in presence of 5 μ M A23187. The defect in ΔPRP1 parasites therefore appears to completely defective in their ability to respond to high spikes in Ca^2+^ concentration with microneme secretion.

The modest reduction in MIC2 secretion in absence of PRP1 using ethanol Fig 3A is not unsurprising since ethanol is not as powerful as the other secretagogues. Thus, this might highlight more subtle effects, especially since it acts at the bifurcation between Ca^2+^ and PA pathways. The titration of ethanol does not reveal a detectable difference in sensitivity toward MIC2 secretion, however, it appeared in some experiments that the total MIC2 amount in ΔPRP1 parasites is slightly lower than in the control line (Fig S3). To determine whether the overall MIC2 content was different in ΔPRP1 parasites we probed a serial dilution series of total parasite lysate with MIC2 antibody and two loading controls, dense granule protein GRA1 and cytoskeleton protein IMC1 (Fig 3C). These data show that the overall MIC2 level remains unchanged upon loss of PRP1. Since PFUS in *Paramecium* has been show to affect secretory organelle formation (Liu et al., 2011), we reasoned that the possible mis-trafficking of the microneme proteins in the ΔPRP1 mutants might also explain the secretion defect. We tested this, as well as trafficking to the other secretory organelles, by fluorescence imaging using MIC2 antibody alongside rhoptry and dense granule specific antisera as controls (Fig 3D). We observed no accumulations of MIC2 protein along the secretory pathway and the morphology of all secretory organelles, including the micronemes, was normal. Thus, the loss of PRP1 does not affect organellogenesis or protein trafficking to the micronemes, but results in the inability to enhance microneme secretion in response to high Ca^2+^ spikes.

### Where does PRP1 localize?

PRP1 was previously shown to localize to the most apical micronemes, and to transition to the cytoplasm upon triggering microneme secretion with ethanol (Matthiesen et al., 2001, Matthiesen et al., 2003). We used our specific PRP1 antiserum in IFA analysis to confirm this observation. Using previously reported 4% paraformaldehyde (PFA) fixation we did not observe any specific signal as the random spotty αPRP1 pattern in wild type parasites looks identical to the signal observed in the ΔPRP1 line (Fig 4A). This suggests that PFA fixation destroys the epitope(s) recognized by the antiserum. Presence of a single band of approximately 64 kDa, which is around the predicted size of 70 kDa, only in wild type parasites clearly indicates the high specificity of our antiserum (Fig 2B). To overcome this fixation artefact we used low temperature 100% methanol fixation and observed cytoplasmic localization of PRP1, both in intracellular and extracellular parasites (Fig 4A). Importantly, we did not observe any significant signal in ΔPRP1 parasites under these conditions except for the background foci also evident with 4% PFA. Taken together, our antiserum is specific for PRP1 but the epitope(s) it recognizes are not preserved upon PFA fixation, which complicates direct comparison to previous reports (Matthiesen et al., 2001, Matthiesen et al., 2003).

**Figure 4.**
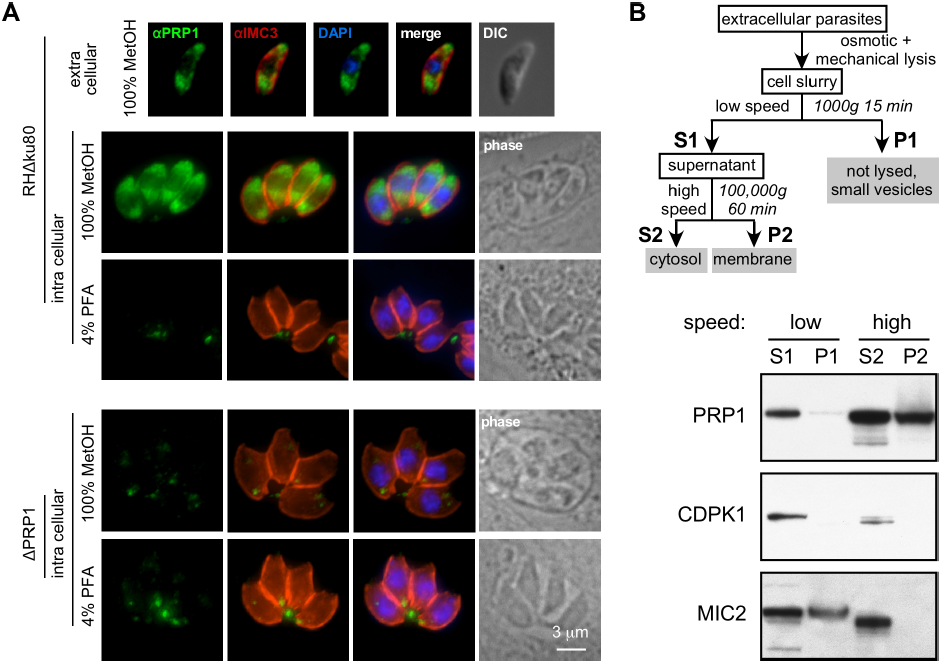
Sub-cellular localization of PRP1 by antiserum. **A.** PRP1 antiserum was used to probe methanol (MetOH) or 4% paraformaldehyde (PFA) fixed wild type (RHΔku80) and ΔPRP1 parasites, intracellular or extracellular, as indicated. IMC3 antiserum was used as control for the cortical cytoskeleton; DAPI stains DNA. Note that PFA fixation either destroys the epitope recognized by the PRP1 antiserum or destroys the structure to which PRP1 localizes. **B.**Fractionation of wild type parasites shows that PRP1 is present in both a membrane associated and non-membrane associated, soluble fraction. Top: flowchart of fractionation strategy. Bottom: western blots of fractions probed with antisera as indicated. Protein sizes as follows: PRP1, 64 kDa; CDPK1, 65 kDa; MIC2, 87 kDa.

To further elucidate the conflicting PRP1 localizations we tagged the endogenous locus at the C-terminus with a yellow fluorescent protein (gPRP1-YFP; Fig 5A, B). Live parasite imaging revealed a cortical YFP signal in combination with an extended apical spot in intracellular parasites, the latter of which was reduced in extracellular parasites (Fig 5C). Co-transfections targeting red fluorescent fusion proteins to the cortical IMC cytoskeleton (IMC1), rhoptries (TLN1) and micronemes (MIC8) showed that the cortical gPRP1-YFP signal colocalized with the IMC whereas the elongated apical signal in intracellular parasites overlays with the rhoptries (Fig 5D). Surprisingly, we observed no co-localization with the micronemes under any condition tested. We further used this YFP tagged PRP1 line to validate our αPRP1 antiserum upon methanol and PFA fixation through co-staining with an αGFP antiserum recognizing YFP. Consolidating our observations made in Fig 4, we observed a cytoplasmic PRP1 and GFP signal upon methanol fixation, but upon PFA fixation we loose the cytoplasmic PRP1 signal, which becomes spotty and does not overlay with the even GFP signal. Thus, PFA fixation indeed destroys the PRP1 epitope(s) recognized by our antiserum. Furthermore, methanol fixation disrupts the PRP1-YFP localization pattern observed in live parasites (Fig 5E). Since membrane associated proteins are notoriously challenging to fixate (e.g. (Hannah *et al.*, 1998)) we explored a variety of fixation conditions to test if the live localization could be conserved (Fig S4). Acetone fixation preserved the cortical signal to some extent, but the rhoptry signal is not preserved under any of the tested conditions. Finally, we wanted to ascertain that tagging the endogenous locus with YFP did not interfere with PRP1 function. Therefore we performed the microneme secretion assay on the gPRP1-YFP line and compared it directly with the parent line (RHΔku80) and the ΔPRP1 line. We detect secreted MIC2 in the YFP tagged line comparable to the parent line, which indicates that PRP1 function is not impaired by the YFP tag (Fig 5F). Overall, our data indicate that all tested fixation conditions alter the PRP1 localization in the cell and that PFA destroys the epitope(s) recognized by our antiserum. We conclude that endogenously tagged PRP1 is functional and localizes to the IMC and rhoptries in intracellular parasites whereas the rhoptry signal is diminished in extracellular parasites. When we stimulated extracellular parasites with A23817 no further changes in localization were observed but we noted extrusion of the conoid suggesting other Ca^2+^-mediated events are executed normally (data not shown).

**Figure 5.**
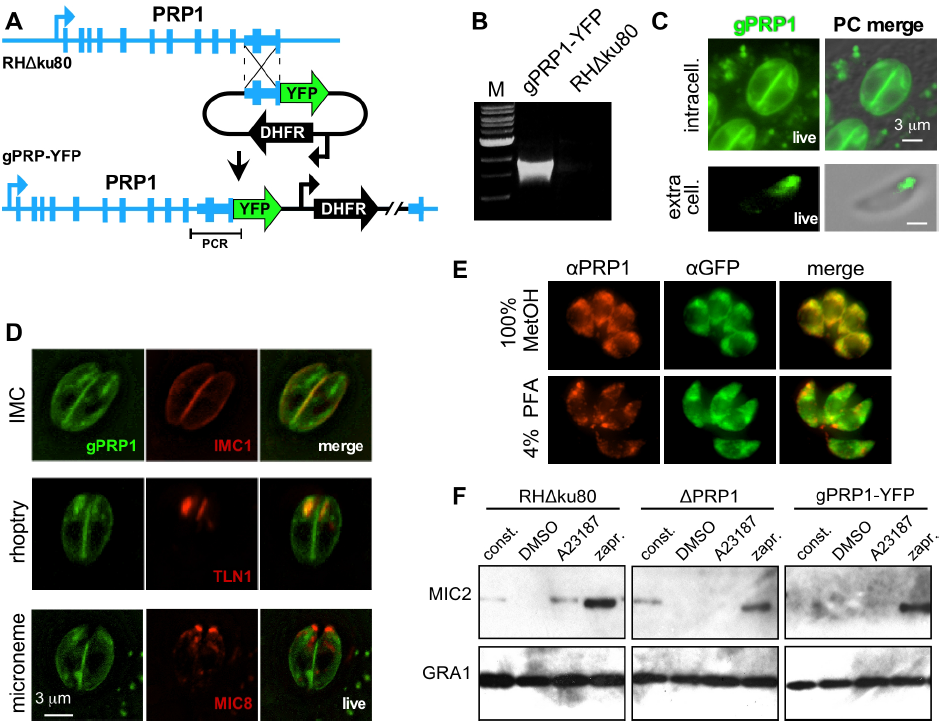
Sub-cellular localization of PRP1 through endogenous tagging. **A.** Schematic representation of generating of C-terminal endogenously YFP-tagged gPRP1-YFP parasites by single homologous recombination into the RHΔku80 parent line. **B.** PCR validation of gPRP1-YFP genotype using the primer pair shown in panel A. **C.** Live imaging of gPRP1-YFP parasites under intracellular and extracellular conditions as indicated. PC: phase-contrast. **D.** Live imaging of gPRP1-YFP parasites co-transfected with markers for the IMC (IMC1-mCherry), rhoptries (TLN1-mCherry), and micronemes (MIC8-mCherry). **E.** Representative images of intracellular gPRP1-YFP parasites fixed using either 100% methanol (MetOH) or 4% paraformaldehyde (PFA) stained with α-PRP1 and α-GFP antisera as indicated. Note that PFA fixation destroys the co-staining of GFP and PRP1 and thus destroys the PRP1 epitope(s) recognized by the specific anti-serum. **F.** Representative western blot of microneme secretion assay on gPRP1-YFP parasites and RHΔku80 and ΔPRP1 controls. Extracellular tachyzoites were treated with A23187 (A), zaprinast (zapr.) or DMSO (D; mock) for five minutes at 37°C. For constitutive (const.) secretion, the tachyzoites were allowed to release the protein for one hour. Dense granule protein GRA1 was used as control for microneme- and Ca^2+^-independent secretion.

Since our live and fixed localizations both conflict with the previously reported microneme localization we further corroborated the membranous localization in live parasites by fractionating wild type parasites in membrane and cytosolic fractions by differential centrifugation (Fig 4B). We used CDPK1 as a cytosolic marker (Pomel *et al.*, 2008) and MIC2 antiserum to probe for micronemal protein. These data show that PRP1 is present as both a soluble and membrane associated form but does not appear to co-fractionate with the micronemes (P1 in Fig 4B). Thus the fractionation data are consistent with the live microscopy observations of the endogenously YFP-tagged PRP1 protein.

### PGM2 also functions in microneme secretion

To ensure that the phenotypic effects observed in the ΔPRP1 mutants are specific to PRP1 deletion and not due to potential functional compensation by the PGM2 paralog we also deleted this gene. We replaced the *pgm2* locus with a drug resistance cassette mediated by a CRISPR/Cas9 induced double strand break, both in parent (RHΔku80) and ΔPRP1 parasites (Fig 6A-C). Plaque assays showed that both the single (ΔPRP1 and ΔPGM2) as well as the double (ΔPRP1ΔPGM2) knock out lines were viable and did not show a dramatic change in plaque size or number (Fig 6D). To specifically understand the effect of deleting PGM2 and determine its role in microneme secretion, we performed secretion assays similar to those with ΔPRP1 and the parental strain (Fig 6E). Surprisingly, both single deletion of PGM2 as well as double knockout of PRP1 and PGM2 demonstrated microneme secretion defects as observed upon absence of only the PRP1 gene: A23187 induced MIC2 secretion was either below or just at the detection limit in all the three mutants whereas there was a modest reduction in ethanol induced secretion. As for ΔPGM2, constitutive MIC2 release or release upon zaprinast and propranolol triggers was unchanged and comparable to the parent line. Thus, our data suggest that both PRP1 and PGM2 act in the microneme secretion pathway, thereby eliminating putatively compensatory functions between the two proteins.

**Figure 6.**
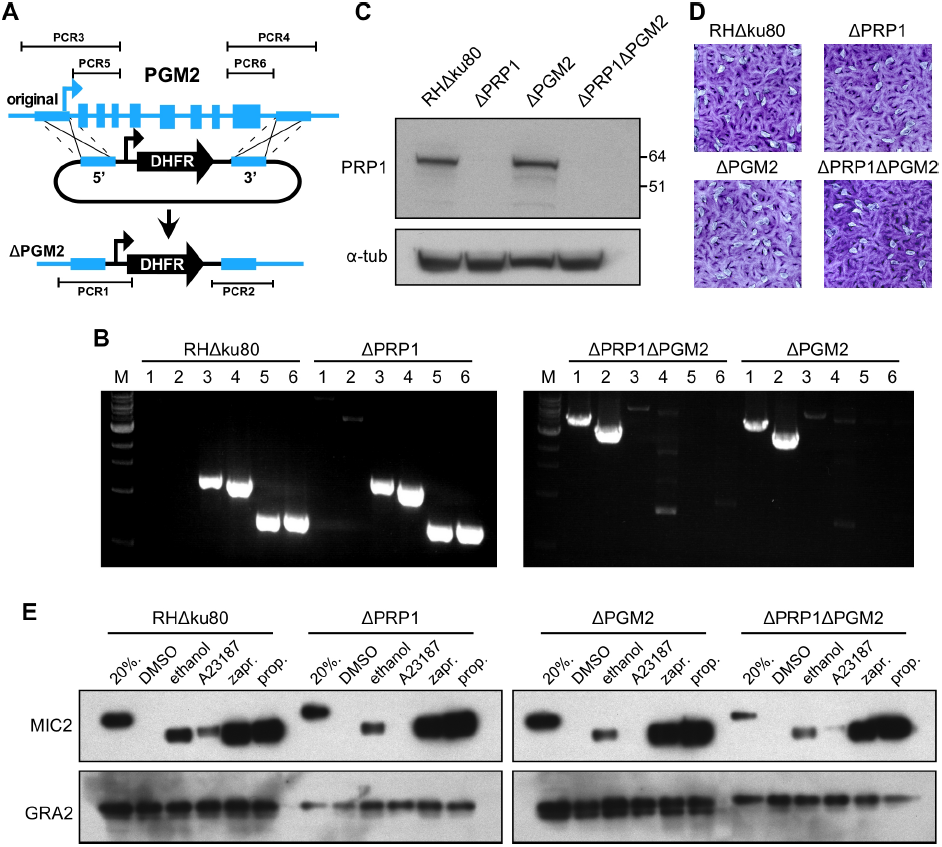
PGM2 is also required for Ca^2+^-trigger induced microneme secretion and does not complement for loss of PRP1. **A.** Schematic representation of generating PGM2 knockouts by double homologous recombination into the RHΔku80 or ΔPRP1 as parent lines. Blue boxes indicate the homologous regions used to replace the endogenous locus. The light blue boxes indicate the exons in the genomic locus of *pgm2*. PCR 1-5 indicate the diagnostic PCR reactions (shown in B). B. Diagnostic PCR reactions validating the replacement of the PGM2 locus with the DHFR cassette in both the RHΔku80 and ΔPRP1 background. M represents 1 kb DNA ladder (NEB). **C.** Diagnostic western blot to validate the presence or absence of PRP1 protein in the generated ΔPGM2 parasite lines in the RHΔku80 or ΔPRP1 backgrounds, respectively. α-tubulin antibody was used as loading control. Numbers in the left represent the protein molecular weight marker. **D.** Representative images of plaque assays performed with parasite lines as indicated, grown undisturbed on a host cell monolayer for seven days. **E.** Representative western blot image of the MIC2 microneme secretion assay. 20% represents the non-secreted total protein from 20% of the total parasites used for the secretion assay. Extracellular tachyzoites were treated with DMSO control, ethanol, A23187, zaprinast (zapr.), or propranolol (prop.) for 5 min at 37°C or assayed for constitutive (const.) secretion for 1 hr. Dense granule protein GRA2 was used as control for microneme- and Ca^2+^-independent secretion.

The shared phenotype between the ΔPRP1 and ΔPGM2 suggest that PRP1 and PGM2 interact functionally. Extending on this thought, they might associate with each other. Proteins within the alpha-D-phosphohexomutase superfamily to which the PGMs belong in general form dimers (e.g. *Paramecium* parafusin is a symmetric dimer: (Muller *et al.*, 2002)), whereas there is an example of a heterodimer of allelic variants (Andreotti *et al.*, 2015). As such we explored the possibility of PRP1/PGM2 heterodimers in *Toxoplasma* tachyzoites. To this end we generated a parasite line expressing a tandem Myc epitope fusion of PGM2 (Myc2-PGM2) and performed a co-immunoprecipitation with Myc-recognizing 9E10 antibodies and probed with our PRP1 specific antiserum. We readily pulled down the Myc2-PGM2 in this assay, however we were unable to detect PRP1 in the same fraction (Fig 7). These data suggest that PRP1 and PGM2 do not physically interact with each other.

**Figure 7.**
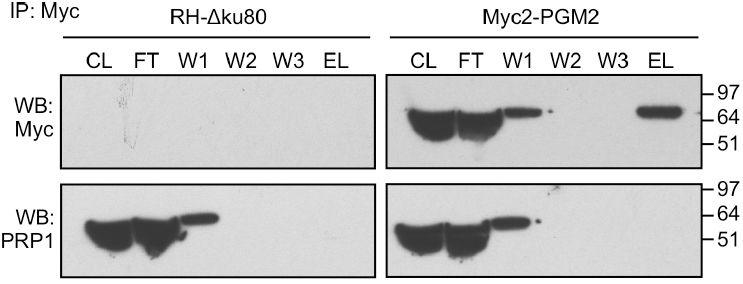
PRP1 and PGM2 do not form a heterodimer. RHΔku80 parasites expressing Myc2-PGM2 or untransfected control parasites were immunoprecipitated with α-Myc beads and western blots probed with either Myc or PRP1 antisera as indicated. No interaction between Myc2-PGM2 and PRP1 is detected. CL: cleared lysate; FT: flow through; W1, 2, 3: wash 1, 2, 3; EL: eluate.

### Neither PRP1 nor PGM2 are required for energy metabolism and acute virulence

Besides the effect on microneme secretion it came somewhat as a surprise that the PRP1 and PGM2 double KO strains were was viable, without an apparent effect on glycolysis (Fig 6D). Furthermore, a direct *in vitro* comparison of the enzymatic efficiency of both paralogs indicated that PRP displayed a 4-fold greater enzymatic activity than PGM2 (Imada et al., 2010), which is also counter intuitive. Although the dispensability of essential glycolytic enzymes in *Toxoplasma* has been reported, such parasites have significant fitness defects and typically require glutamine complementation (Shen & Sibley, 2014, MacRae *et al.*, 2012, Blume *et al.*, 2009, Dubey *et al.*, 2016, Nitzsche *et al.*, 2016). However, the knock down of PGM in *Saccharomyces cerevisiae* (Fu et al., 2000) or depletion of the single PGM (PFUS) in *Paramecium* and *Tetrahymena* (Liu et al., 2011, Chilcoat & Turkewitz, 1997) demonstrated no apparent effect on the growth of these organisms, and there are organisms without a dedicated PGM such as *Trypanosoma brucei* (Bandini *et al.*, 2012). In addition, we showed that the PFUS-related PGM orthologs vs. the exclusive PGM enzymes are not at all uniformly conserved across the Apicomplexa (Fig 1), which further supporting the notion that these proteins display plasticity in these organisms. Therefore, we asked if loss of either one or both PGM orthologs in *Toxoplasma* had an effect on metabolism by assessing the ATP level in the knock-out mutants we generated. As shown in Fig 8A, no changes in ATP levels were observed in the different mutants compared to the wild type parent line. These data suggest that the *Toxoplasma* metabolism shares characteristics with the organisms where PGM is dispensable and we conclude that PGM activity is not required for the *Toxoplasma* lytic cycle.

**Figure 8.**
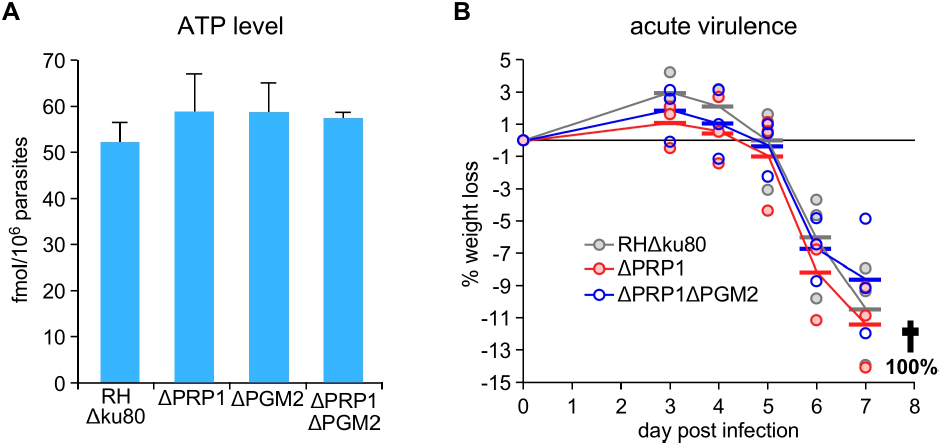
Loss of the PRP1/PGM2 complex does not affect metabolism or acute virulence in mice. **A.** ATP levels were determined for intracellularly growing parasite lines as indicated. n=3 +s.d. B. Acute virulence in BALB/c mice was assessed by i.p. infection of 1,000 tachyzoites of the parasite lines as indicated. Weight changes relative to day 0 are shown for each mouse; averages per group are show by the horizontal bar; plotted lines represent average. All mice in each group either died or were euthanized at day 8 p.i.

Across the metabolic and Ca^2+^-dependent exocytosis functions of the PGM orthologs in *Toxoplasma* we observe a role in microneme secretion but no effect on plaque forming capacity of tachyzoite *in vitro*. However, it is possible that either function is only prominently required for animal infection, where the environment is much more complex. Germane to this point, Ca^2+^ is known to be critical for virulence and egress to escape the innate immune response during acute mice infection (Tomita *et al.*, 2009). It is conceivable that this escape response needs to be fast and could rely on high spikes of intracellular Ca^2+^. To test this model we assessed the acute virulence of the ΔPRP1 and ΔPRP1ΔPGM2 mutants through mice infections. We observed no difference in the development of the infections between either mutant or compared to the RHΔku80 parent line (Fig 8B). Weight loss patterns were comparable for all infected mice and all mice either died or needed euthanasia on day 8 post infection. Thus, these data indicate that the PGM orthologs are not only dispensable *in vitro*, but also *in vivo* during an acute infection.

## Discussion

Ca^2+^-mediated exocytosis is a key event required for successful host cell invasion of apicomplexan parasites. *Toxoplasma* PRP1 was associated with this process based on localization studies in *Toxoplasma* and heterologous functional studies in the ciliate *Paramecium* (Matthiesen et al., 2003). Here we directly assessed the role of PRP1 in *Toxoplasma* by deleting the gene from the parasite, which surprisingly resulted in a non-lethal phenotype both *in vitro* and *in vivo* (Fig 2, 8B). We did, however identify a defect in the release of micronemes triggered by elevated Ca^2+^ concentrations. Interestingly, our detailed analysis of the PRP1 deletion mutant revealed that the morphology of the secretory organelles and cytoskeletal inner membrane complex appear normal in the ΔPRP1 line (Fig 3D, 4A). Furthermore, we demonstrated that invasion and egress efficiency, which are both reliant on successful Ca^2+^-mediated exocytosis, are independent of PRP1 (Fig 2C,D).

Since PRP1 was hypothesized to play a role in microneme secretion we determined the dynamics of *in vitro* microneme secretion under different conditions and stimulations. Our work demonstrates for the first time that deletion of PRP1 only affects microneme secretion bursts triggered by a high Ca^2+^ concentration through stimulation with the Ca^2+^-ionophore A23187 (Fig 3A). Titration of A23187 demonstrated that the loss of PRP1 completely disrupts the secretion boost in response to high Ca^2+^ and is not lowering the amplification of this high Ca^2+^ secretion response by lowering the general sensitivity to Ca^2+^ (Fig 3B). Overall, these data indicate that PRP1 is essential to translate a high Ca^2+^ trigger into enhanced microneme secretion. Spikes in intracellular Ca^2+^ are known to occur during invasion and egress (Sidik et al., 2016a, Wetzel et al., 2004), which correlate with increased levels of microneme secretion. However, since we observe no fitness loss of the ΔPRP1 line the physiological function of these spikes in microneme secretion is currently not clear. ΔPRP1 have microneme secretion levels compared to the wild type under constitutive secretion in the extracellular environment, in absence of pharmacological stimuli. Our data therefore suggest that a basal level of microneme secretion is sufficient to complete the lytic cycle of the parasite. It is possible that the absence of PRP1 has certain cell biological effects such as changes in gliding motility patterns as observed in other mutants. In addition, we cannot exclude that the ability to respond to high Ca^2+^ fluxes with a microneme secretion response might be critical during other developmental stages of the parasite.

The PRP1 mutant allowed us to dissect the contribution of different branches of the signal transduction to microneme secretion through stimulation with targeted pharmacological secretagogues. Upstream stimulation of the pathway with PKG-activating zaprinast leads to robust microneme secretion. In *Plasmodium* numerous protein substrates have been described as PKG substrates (Brochet *et al.*, 2014), which likely have many parallels in *Toxoplasma*. In addition, PKG activity generates phosphatidylinositol 4,5-bisphosphate (PI_(4,5)_P_2_) through the phosphorylation of phosphatidylinositol (PI) and was shown to be a key event in *Plasmodium* secretion (Brochet et al., 2014). PI_(4,5)_P_2_ is the substrate for PI-PLC, which in turn forms IP_3_ and DAG. IP_3_ is associated with inducing the Ca^2+^ spike resulting in microneme secretion (Lovett *et al.*, 2002, Sidik *et al.*, 2016b), whereas DAG is converted in PA, which is directly sensed by the micronemes to promote their secretion (Bullen et al., 2016). In contrast to PKG stimulation, PI-PLC activation by ethanol results in mildly reduced secretion in absence of PRP1 (Fig 3A). These results show that the phosphorylation of PI by PKG is a major, but not the only requirement for a complete microneme secretion response, thereby implying the presence of other contributing PKG substrates. Excluding the contribution of the Ca^2+^-mediated branch by specifically triggering PA production through propranolol stimulation releases all brakes on microneme secretion (Bullen et al., 2016) and leads to a higher secretion than PKG stimulation regardless of the presence of PRP1. This indicates that very high PA levels override the requirement of substrates other than PI for PKG. Thus, our work provides evidence for PKG substrates other than PI contributing to the microneme secretion response. In ciliates PFUS is phosphorylated by casein kinase II (CKII) and PKG (Kussmann *et al.*, 1999) while during exocytosis the protein undergoes dephosphorylation by the protein phosphatase calcineurin (CN) (Plattner & Kissmehl, 2005, Treptau *et al.*, 1995, Subramanian & Satir, 1992, Kissmehl *et al.*, 1996). The dephosphorylation results in dissociation of PFUS from the ciliate secreting vesicles while the released PFUS re-associates with nascent vesicles upon phosphorylation. It is conceivable that PRP1 phosphorylation by PKG is required for microneme secretion, but our data on *Toxoplasma* CN demonstrate this is not the key activity of CN as microneme secretion remains unaffected upon CN depletion (Paul *et al.*, 2015). The *Toxoplasma* phosphoproteome validated two phosphorylation sites in PRP1, Thr156 and Ser158, and the phosphorylation of the orthologous residues Thr156 and Ser158 in PGM2. However, the phosphorylation of these residues is universal requirement for enzymatic activity in phosphohexomutase family proteins (Bandini et al., 2012), and hence are unlikely to be connected to regulation of Ca^2+^-dependent exocytosis. Amongst the other detected phosphorylation sites in PFUS with reasonable confidence, four are conserved in PRP1.Additional work is needed to establish a possible connection of these putative phosphorylation with exocytosis.

Through the use of a peptide antibody against PFUS, which cross-reacts with PRP1, the latter was reported to localize to the most apical micronemes from which it releases in a phosphorylation state dependent fashion upon secretion (Matthiesen et al., 2001). In this model, PRP1 prevents simultaneous secretion of all micronemes in a parasite, which would be detrimental as a dosed release of micronemes is required to support prolonged gliding for crossing biological barriers while reserving micronemes needed for successful host cell invasion. However, the observed secretion dynamics of the PRP1 deletion mutant are inconsistent with this model. Moreover, our localization data also conflict with these previous results as the endogenously fused YFP associates with the IMC and rhoptries (Fig 5). Although highly specific in western blots, unfortunately our PRP1 antiserum was incompatible with IFA fixation methods, thereby restraining us from performing the same experiments (Fig 2, 4B, 7). Membrane associated proteins are notoriously challenging to localize sub-cellularly due to fixation artefacts (Hannah et al., 1998) which is mirrored by our experience. However, we were able to confirm that PRP1 is present in soluble and membrane associated forms in extracellular parasites, which is consistent with the previously presented model (Liu *et al.*, 2009). In our opinion, the YFP-fusion data are the best reflection of the natural PRP1 localization, however, how the IMC and rhoptry localization of PRP1 relate to its role in microneme secretion is presently unclear.

To ensure that the phenotypic effects observed in the ΔPRP1 mutants are specific to PRP1 deletion and not due to potential functional compensation by the PGM2 paralog we also removed this gene and even made a double knock out mutant (Fig 6). The observation that deletion of either one or both PGM genes simultaneously results in the same microneme secretion defect upon high Ca^2+^ flux sheds a completely new light on the specific role of PRP1 in Ca^2+^ signaling: PRP1 is not exclusive in this function as it extends to the other PGM expressed in *Toxoplasma*. This observation might explain why the conservation of either ortholog in the various Apicomplexa is a mishmash (Fig 1) as either ortholog could function in Ca^2+^-dependent secretion. However, that individual gene deletions in *Toxoplasma* resulted in the same phenotype suggests a functional interaction between PRP1 and PGM2, which suggest a likely selective evolutionary pressure on preserving both. By co-IP we did not detect a physical interaction (Fig 7). Alternatively, the interaction might be transient upon high Ca^2+^, which is much harder to capture.

In addition, our data show that *Toxoplasma* tachyzoites can survive without dedicated phosphoglucomutase. Since PGMs function in glycolysis this absence was expected to affect energy production, however we did not detect a change in ATP level in parasites without PGM (Fig 8A). Similar observations have been made in yeast (Hofmann *et al.*, 1994, Boles *et al.*, 1994). Moreover, other organisms such as *Trypanosoma brucei* do not have a dedicated PGM enzyme at all (Bandini et al., 2012). In these and other systems it has been established that related enzymes in the alpha-D-phosphohexomutase superfamily also have phosphoglucomutase activity, such as phosphomannomutase (PMM) and phopho N-acetylglucosaminemutase (PAGM). Indeed *Toxoplasma* encodes a PMM (TGME49_239710) and a PAGM (TGME49_264650) and the amino acids involved in sugar binding are conserved between the *Toxoplasma* and the *T. brucei* PMM (Bandini et al., 2012). These insights therefore explain why the deletion of both *Toxoplasma* PGMs has no effect on energy production.

In conclusion, we show that PRP1 is partially membrane associated and functions in translating a spike in Ca^2+^ into a burst of microneme secretion. The PRP1 line also confirmed the bifurcation in the Ca^2+^ and PA mediated control of microneme secretion downstream of PI-PLC activity. The constitutive level of microneme secretion is independent of PRP1 and sufficient to support the parasite through all steps of the lytic cycle both *in vitro* and *in vivo*. Interestingly, neither *Toxoplasma* PGM paralog is required for glycolysis whereas deletion of either gene results in loss of the microneme secretion burst. Considering our data in context of the mosaic pattern of PGM ortholog conservation across the Apicomplexa, our data suggests that either PGM gene could function in controlling the secretory response upon spikes in Ca^2+^, which indicates this control mechanism could be more wide spread than initially suspected.

## Material and Methods

### Parasites and host cells

*Toxoplasma* tachyzoites were maintained in human foreskin fibroblasts (HFF) as previously described (Roos *et al.*, 1994). The host cells were maintained in DMEM media containing 10% serum. *Toxoplasma* strain RHΔku80ΔHX (Huynh & Carruthers, 2009) was used as the basis for all mutants in this study. Stable transgenics were obtained by selection under 1 μ M pyrimethamine, 25 μ g/ml mycophenolic acid (MPA) combined with 50 μ g/ml xanthine or 20μ M chloramphenicol and cloned by limiting dilution.

### Plasmids

All primer sequences are provided in supplementary table S1. For tagging the endogenous PRP1, 1.5 kb genomic DNA upstream of stop codon was PCR amplified using primer pair PRP1-LIC-F/R and cloned by ligation independent cloning into plasmid pYFP-LIC-DHFR (kindly provided by Vern Carruthers, University of Michigan) (Huynh & Carruthers, 2009). The plasmid was linearized with *Nco*I prior transfection.

For the generation of ΔPRP1 plasmid, we deleted the LoxP flanked tubulin protemer and YFP cassette from the p5RT70-loxP-KillerRed-loxP-YFP/HX plasmid ((Andenmatten *et al.*, 2012), kindly provided by Markus Meissner, University of Glasgow) by *Apa*I/*Not*I digestion, blunting the ends and re-ligation. The 2 kb flanks for double homologous recombination (HR) were PCR amplified from RHΔHX genomic DNA using primer pairs 5‘PRP1-KpnI-F/R for the 5’ flank located 1.3 kb upstream of the ATG start codon and 3‘PRP-SacI-F/R for the 3’ flank downstream of translation stop. The 5‘-flank was inserted into p5RT70--/HX using *Kpn*I and independently the 3’flank was inserted separately in parallel into p5RT70--/HX using *Sac*I; the two plasmids were conjoined using *Sca*I/*Not*I to make the final double homologous recombination plasmid. The plasmid was linearized with *Sca*I prior to transfection.

Plasmid ptub-Myc2-PGM2/sagCAT was cloned by PCR amplifying the PGM2 CDS from cDNA using primer pair PGM2-AvrII-F and PGM2-EcoRV-R and was cloned by *Avr*II/*Eco*RV digestion into the ptub-Myc2-GAPDH1/CAT (Dubey et al., 2016).

### Generation of PGM2 knockout

To generate ΔPGM2 parasite, two CRISPR/Cas9 plasmids targeting Cas9 to the *pgm2* genomic locus were designed using the primer pairs 5PGM2-dKO-s/as and 3PGM2-dKO-s/as respectively upstream of AUG and downstream of stop codon. Specificity of the guide RNA was tested as previously published (Sidik *et al.*, 2014). DHFR drug selectable cassette along with 20 nt of homologous region flanking each end which corresponds to upstream and downstream of the genomic locus was PCR amplified. 40μ g of each pU6-5’PGM2/3’PGM2-Cas9 plasmid was co-transfected with 40 μ g of the PCR product into RHΔKu80ΔHX or RHΔKu80ΔHXΔPRP1 parasites to generate ΔPGM2.

### Immunofluorescence assays and live fluorescence microscopy

Live microscopy was performed on intracellular parasites grown overnight in 6-well plate containing coverslips confluent with were filtered, pelleted and resuspended in PBS. Thereafter, parasites were added to poly-L-lysine coated cover-slips and allowed to incubate for 30 minutes at 4° C prior to imaging. Co-localization studies in live gPRP1-YFP parasites was performed following transient co-transfection with the following plasmids: tub-IMC1mCherryRFP/sagCAT (Anderson-White *et al.*, 2011); TLN1(1-58)mCherryRFP/HPT (rhoptry marker) (Hajagos *et al.*, 2012) (kindly provided by Peter Bradley, UCLA); pTgMIC8-TgMIC8mycmCherryFP-nosel (pG53) (Kessler *et al.*, 2008) (kindly provided by Markus Meissner, University of Glasgow).

Indirect immunofluorescence assay was performed on intracellular parasites grown overnight in 6-well plate containing coverslips confluent with HFF cells fixed with 100% methanol (unless stated otherwise) using the following primary antisera: rabbit αGFP (1:500) (Torrey Pines Biolabs), α-MIC2 MAb 6D10 (1:8,000; kindly provided by David Sibley, Washington University in St. Louis (Wan *et al.*, 1997)), rabbit α-MIC2 (1:8,000; kindly provided by David Sibley, Washington University in St. Louis) (Labruyere *et al.*, 1999), mouse α-ROP1 (1:1,000; kindly provided by Peter Bradley, UCLA (Saffer *et al.*, 1992)), α-GRA1 (1:20,000; kindly provided by Marie-France Cesbron-Delauw, Université Grenoble, France (Cesbron-Delauw *et al.*, 1989)), rat α-IMC3 (1:2,000 (Anderson-White et al., 2011)), and guinea pig α-PRP1 (1:1,000). Alexa 488 (A488) or A594 conjugated goat α-mouse, α-rabbit, α-rat, or α-guinea pig were used as secondary antibodies (1:500, Invitrogen). DNA was stained with 4’,6-diamidino-2-phenylindole (DAPI). Fixation and staining of P-STAT3 (1:120; #9145P Cell Signaling) was performed exactly as described in (Brown *et al.*, 2014). A Zeiss Axiovert 200 M wide-field fluorescence microscope equipped with a α-Plan-Fluar 100x/1.3 NA and 100x/1.45 NA oil objectives and a Hamamatsu C4742-95 CCD camera was used to collect images, which were deconvolved and adjusted for phase contrast using Volocity software (Improvision/Perkin Elmer).

For testing the different fixatives and the fixation conditions for PRP1 localization, 100% acetone; 4% paraformaldehyde (PFA); 100% methanol and 2.5% formaldehyde and 0.05% glutaraldehyde at −20°C, 4°C or room temperature were tested.

### Generation of specific PRP1 antiserum

To generate N-terminal His_6_ tagged fusion protein, 580 bp from the cDNA of PRP1 (corresponding to amino acids: 446 to 637 in the C-terminal region of the protein) were PCR amplified using the primers aPRP1-LIC-F/R and cloned into the pAVA0421 plasmid using LIC (Alexandrov *et al.*, 2004). The fusion protein was expressed in BL21 *Escherichia coli* using 1 mM IPTG for overnight at 37° C and purified under denaturing condition over Ni-NTA Agarose (Invitrogen). Polyclonal antiserum was generated by immunization of guinea pig (Covance, Denver, PA).

### Western blotting

Following SDS PAGE, PVDF membrane blots were probed with mouse α-IMC1 (1:2,000, kindly provided by Gary Ward, University of Vermont), guinea pig α-PRP1 (1:10,000), mouse monoclonal α-GRA1 (1:20,000), or CDPK1 nanobody (1 μg/ml, a kind gift of Dr.Sebastian Lourido, Whitehead Institute (Ingram *et al.*, 2015)) followed by probing with horse radish peroxidase (HRP)-conjugated α-mouse (1:10,000), α-guinea-pig antibody (1:3,000, Santa-Cruz Biotech), or in case of the nanobody, α-penta-His antibody (1:10,000; Qiagen) and detection of signal by chemiluminescent HRP substrate (Millipore).

### Fractionation

Sub-cellular fractionation was performed as described previously (Dubey et al., 2016). Essentially, extracellular parasites were lysed by 5 min freezing of the pellet in liquid nitrogen and resuspension at 37° C in hypotonic buffer (10 mM Tris pH 7.8 and 5 mM NaCl) followed by additional lyses by 40 douncer strokes. The lysate was centrifuged at low speed (1000*g* for 15 min). The pellet was resuspended in an equal volume of resuspension buffer (100 mM Tris pH 7.8 and 150 mM NaCl) and centrifuged at high speed (100,000*g* for 60 min). For SDS PAGE pellets were resuspended in an equal volume of SDS-PAGE loading buffer and corresponding amounts analyzed by western blotting.

### Plaque assay

T12.5 culture flasks or 6-well plates confluent with HFF cells were inoculated with 100 parasites of choice and grown for 7 days. Following incubation, the monolayer was fixed with 100% ethanol for 10 min and stained with crystal violet (Roos et al., 1994). Plaque sizes were quantitated using ImageJ-win32 software (Abramoff, 2004).

### Microneme secretion assay

Microneme secretion assays using the MIC2 was performed as published (Carruthers & Sibley, 1999). Briefly, freshly lysed parasites were pelleted, washed and resuspended to 2× 10^8^/ml parasites in DMEM/FBS (DMEM containing 20 mM HEPES, pH 7.0 and 3% (w/v) FBS). 2× 10^7^ parasites were added to each well of a 96-well plate and secretion was induced by the following secretagogues: 1.25 μ M A23187, 0.25%(v/v) ethanol, 500 μM zaprinast, 500 μM propranolol or DMSO control for five minutes at 37° C, unless concentrations are specifically indicated otherwise. For constitutive microneme secretion, we incubated parasites at 37°C for 60 minutes in absence of secretagogue. Microneme secretion was assessed in the supernatant with mouse monoclonal α-MIC2 6D10 (1:8,000). Ca^2+^-independent constitutive secretion of the dense granules was determined using mouse monoclonal α-GRA1 (1:20,000) or rabbit α-GRA2 antiserum (1:10,000).

### Invasion assay

The red-green invasion assay was performed as previously published (Farrell *et al.*, 2012, Kafsack *et al.*, 2004) with modifications. 2.5-3× 10^5^ tachyzoites were added to host cells grown in 96-well black/optical bottom plates, centrifuged (500 rpm, three minutes, RT) and allowed to invade for one hour at 37° C. Non-invading extracellular parasites were detected using A594 conjugated α-SAG1/P30-T41E5 antibodies (1:500; kindly provided by Dr. Jean François Dubremetz, University of Montpellier, France (Couvreur *et al.*, 1988)) or α-SAG1 MAb DG52 (kindly provided by Dr. Jeroen Saeij, Massachusetts Institute of Technology (Burg *et al.*, 1988)). Following 16% formaldehyde/8% glutaraldehyde fixation and permeabilization using 0.25% Triton X-100, the parasites with incubated with A488 conjugated α-SAG1/P30 antibody to visualize both invaded and non-invaded parasites. The antibody incubations were each followed by three washes with HH buffer (Hank‘s Balanced Salt Solution containing 1 mM HEPES, pH 7.0). Cytochalasin D treated wild type parasites were used as negative control. Images were taken using EVOS® FL Cell Imaging System (Life Technologies).

### Egress assay

Egress assay was performed as described previously (Farrell et al., 2012, Eidell *et al.*, 2010). 6-well plates containing coverslips confluent with HFF cells were infected with 6×10^4^ RHΔku80 or ΔPRP1 parasites expressing cytoplasmic YFP (Gubbels *et al.*, 2003) and grown for 30-35 hrs. Egress was triggered by treatment with A23187 or propranolol at concentration as indicated using DMSO as control at 37°C for five minutes, followed by 100% methanol fixation for 10 minutes at RT. Intact vacuoles were counted in at least 10 fields in two independent experiments.

### ATP concentration measurement

The CellTiter-Glo Luminescent Cell Viability Assay (Promega; Madison, WI) was used following the manufacturer’s protocol. Intracellular parasites grown for 18 hrs were harvested and resuspended in phenol red free ED1 medium to 10^7^ parasites/ml. 100 μ l of parasites were added to a 96 well plate and equilibrated and an equal volume of CellTiter-Glo reagent was added. The plate was shaken for 2 min to allow cell lysis and then incubated at room temperature for 30 min prior to luminescence reading on an M5 plate reader (Molecular Dynamics) set at an integration time of 500 ms. An ATP (disodium salt) dilution series was used to generate a standard curve.

### Co-immunoprecipitation

Co-immunoprecipitation basically followed published procedures (Suvorova *et al.*, 2013). Briefly, extracellular parasite pellets were subjected to one freeze-thaw cycle and lysed in lysis buffer (1xPBS, 0.25% NP40, 400 mM NaCl, 250 U/ml benzonase (Novagen), mammalian protease inhibitor cocktail (Sigma)). Lysates were pre-cleared on ProteinG-magnetic beads (New England Biolabs) followed by Myc-tagged protein complex capture on 9E10 monoclonal antibody conjugated magnetic beads (MBL). Beads were washed with three times with lysis buffer and bound proteins eluted in Laemmli buffer.

### *In vivo* mouse infection studies

Groups of three C57BL/6J mice with a weight between 18-20 g were infected intraperitoneally with 1,000 tachyzoites of the RHΔku80, ΔPRP1 or ΔPRP1ΔPGM2 strains. Following infection mice were monitored daily for posture, activity level and weight.

### Sequence analysis and phylogeny

Phylogeny was performed using Geneious (Kearse *et al.*, 2012) and unrooted trees were plotted using the neighbor-joining algorithm.

## Acknowledgements

We thank Dr. Jayme Henzy for assistance with phylogeny and Drs. Giulia Bandini and Sebastian Lourido for insightful discussions. We thank Drs. Peter Bradley, Marie-France Cesbron-Delauw, Jean-François Dubremetz, Sebastian Lourido, Markus Meissner, Jeroen Saeij, David Sibley and Gary Ward for reagents. This work was supported by NIH AI108251 (BIC), AI099658 (MJG), AI122923 (MJG), GM084383 (IJB), AI069986 (IJB) and American Cancer Society RSG-12-175-01-MPC (MJG) grants.

## Additional information

The authors state no competing financial interests.

---

**Supplementary Table S1.**
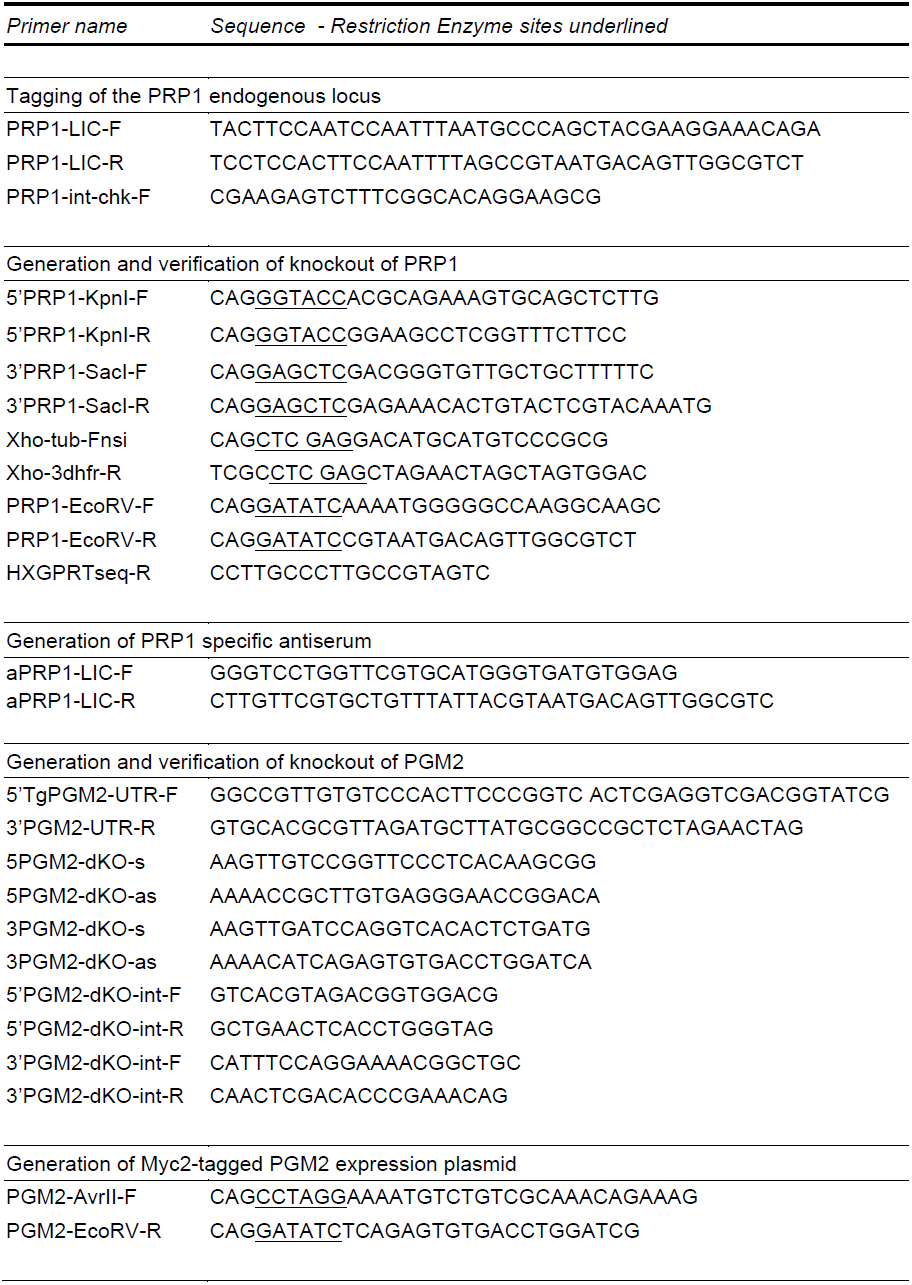
Sequences of primers used.

## Supplementary Figures

**Figure S1.**
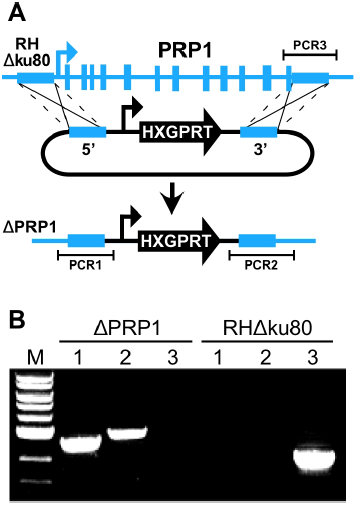
Generation and validation of ΔPRP1 parasite line. **A.** Schematic of the generation of PRP1 knockout parasites. Medium height blue boxes indicate the 5‘ and 3‘ homologous regions used to replace the endogenous locus along with the endogenous promoter; Tall blue boxes represent PRP1 exons. PCR 1, 2 and 3 indicate the amplicons generated by the diagnostic PCRs (shown in panel B). **B.** PCR analysis validating the replacement of the endogenous locus with the drug selection cassette (shown in A) using the genomic DNA of ΔPRP1 and parental RHΔku80 parasites as templates. Specific primer pairs correspond to the PCRs illustrated in panel A. M represents 1 kb DNA ladder (NEB).

**Figure S2.**
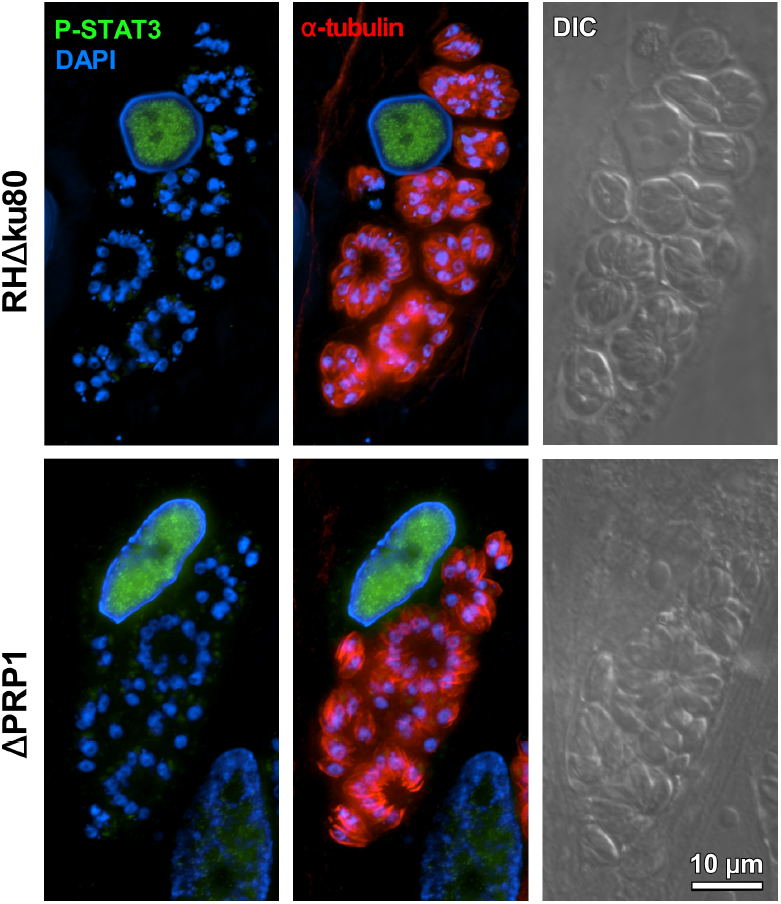
Absence of PRP1 does not interfere with rhoptry secretion. Rhoptry secretion was monitored by phosphorylated STAT3 (P-STAT3) accumulating in in the nucleus of the infected host cell as STAT3 is phosphorylated by the rhoptry protein ROP16 (Saeij et al., 2007). DAPI stains the nuclear material whereas the parasites cytoskeleton is stained with α-tubulin MAb 12G10.

**Figure S3.**
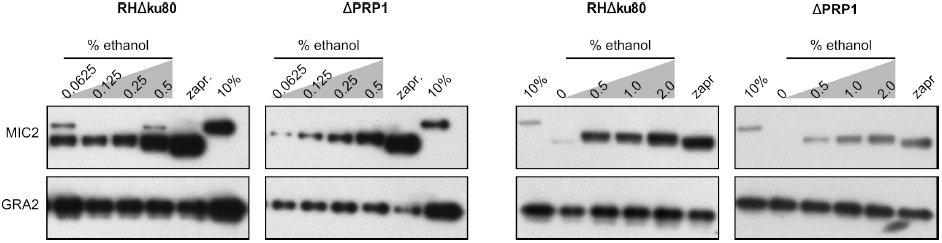
Titration of ethanol induced microneme secretion. Extracellular tachyzoites were treated with different ethanol amounts and zaprinast (zapr.) as indicated for five minutes at 37°C. 10% represents protein lysate from 10% of the total parasites used for the assay. MIC2 was used as the marker for microneme secretion and GRA2 used as the control.

**Figure S4.**
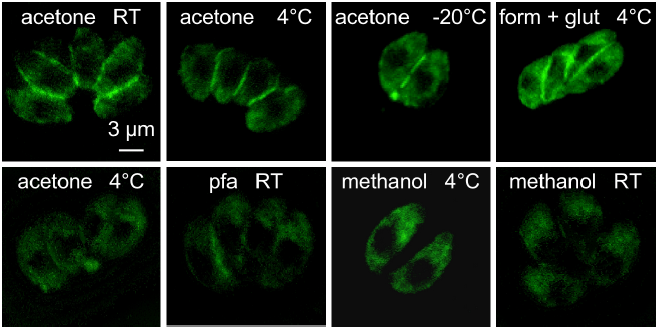
Effect of different fixatives of pPRP1-YFP localization. Representative images of intracellular gPRP1-YFP parasites fixed using different fixative at different temperatures. Acetone: 100% acetone; pfa: 4% paraformaldehyde; methanol: 100% methanol; form + glut: 2.5% formaldehyde and 0.05% glutaraldehyde. RT is room temperature.

